# Performance of the sea turtle *Lepidochelys olivacea* hatchlings from a hatchery on the Pacific coast of Guatemala

**DOI:** 10.1101/2021.09.13.460182

**Authors:** B. Alejandra Morales-Mérida, María Renee Contreras-Mérida, Adriana Cortés-Gómez, Matthew H. Godfrey, Marc Girondot

## Abstract

Sea turtles are marine species that are generally in danger of extinction. The conservation strategies in the different countries are attempting to preserve these species and should be constantly updating their policies according to research results taking place on site. The most abundant and frequent species of sea turtle that nest in the Pacific Coast of Guatemala is *Lepidochelys olivacea* (Eschscholtz, 1829), therefore human predation has been historically high. The solution to this predation, since the 1970s, as a conservation strategy was to place eggs in enclosed protected spaces called hatcheries, where collectors must give 20% of the nest as a conservation quota. Since this program leads to no natural nests (*in situ*) remaining on the beaches, the good functioning of the hatcheries plays a fundamental role in the conservation process to work. To understand and predict the fitness of the hatchlings being produced in Guatemalan hatcheries, crawling performance and self-righting performance were measured in 210 hatchlings of the Multiple Uses Area of Hawaii, in the Pacific Coast of Guatemala. The results of the performance were contrasted with incubation conditions to provide an insight into how the management may influence it. We found that self-righting may be a more meaningful measure of variable behavior than crawling performance, showing that there was little variation due to the homogeneous environment of the hatcheries. We can conclude that a greater number of eggs result in faster self-righting, while deeper nests produce hatchlings with slower rates of self-righting.

**Summary statement:** When hatcheries are the only sea turtle conservation strategy, and their well-functioning is vital to achieve this purpose, performance can provide information of hatchlings’ fitness in response to management conditions.

## Introduction

Temperature plays an important role in the development of sea turtle embryos that incubate inside eggs placed on nesting beaches. In particular, much attention has been focused on the impacts of temperature on hatchling sex ratios, because of temperature-dependent sexual differentiation or TSD (Flores-Aguirre et al., 2020), and on hatching success, because of thermal limits to successful incubation (Howard et al., 2014). Furthermore, projected warming of nesting beaches related to climate change has increased concern about skewed sex ratios and reduced rates of hatching for sea turtles (Hamann et al., 2013, Jensen et al., 2018). In addition to these impacts, incubation temperatures have been shown to impact the behavior and performance of sea turtle hatchlings, which likely has impacts on overall fitness and thus population dynamcis (Booth, 2018).

Performance of hatchlings has been studied to understand and predict the survival of the hatchlings, thus, it can be linked to different factors, depending on the focus of the research (Gatto and Reina, 2020a, Gatto and Reina, 2020b, Booth et al., 2004, Mueller et al., 2019, Booth et al., 2013, Burgess et al., 2006, Sim et al., 2014b, Sim et al., 2014a, Ischer et al., 2009, Booth and Evans, 2011). The most common performance measure studied has been swimming locomotion, in which the activity rate of hatchlings is measured and contrasted with certain factors that are expected to influence the time, such as temperature (Mueller et al., 2019). Another performance factor that has been studied is locomotion, which can be measured by different metrics, including crawling speed, which is measured as the time taken for a hatchling to traverse a set distance, and self righting responses, in which the time required for a hatchling to return to normal position after being placed in a recumbent position (Rivas et al., 2019).

Performance studies in sea turtles have most commonly focused on hatchlings of *Chelonia mydas, Natator depressus*, and *Lepidochelys olivacea* (Gatto and Reina, 2020a, Gatto and Reina, 2020b, Booth et al., 2004, Mueller et al., 2019, Booth et al., 2013, Burgess et al., 2006, Sim et al., 2014b, Sim et al., 2014a, Ischer et al., 2009, Booth and Evans, 2011). In addition, there are some research also done with *Caretta caretta*, and even with freshwater turtles (Booth et al., 2004, Sim et al., 2015, Sim et al., 2014a, Du and Ji, 2003, Elnitsky and Claussen, 2006). Swimming locomotion is assumed to provide the most reliable results because hatchlings engage in swimming immediately after entering the ocean from the beach where were produced. The rate of predation of hatchlings can be high in coastal waters, as they move towards their feeding areas in more open waters (Gyuris 1993). Measurements of swimming performance of sea turtle hatchlings have been typically collected during the first 24 hours after emergence, but various other factors can affect swim speeds, including incubation conditions and light exposure after hatching (Gatto and Reina (2020a).

In addition to looking at impacts of incubation temperature, locomotor and self righting have been measured in *C. mydas, N. depressus*, and *L. olivacea* hatchlings in relation to other incubation factors such as moisture (Gatto and Reina, 2020a). In the latter case, it was shown for *C. mydas*, that moisture had no effect on self righting, while for *L. olivacea* and *N. depressus* hatchlings, increased levels of moisture (from 4-8%) during incubation led to slower self-righting times. In terms of crawl speeds, for *L. olivacea* hatchlings, their crawling was slower at 4% moisture, while there was no relationship between moisture levels of incubation and crawl speeds for *C. mydas* and *N. depressus* hatchlings. It is likely that other factors, including those related to incubation that may influence embryo behavior and performance.

The survival of sea turtle embryos depends on the interaction of several factors during incubation, including salinity, humidity, temperature, gas exchange, rain, storm surge, erosion, and predation (Kaska and Downie, 1999). In locations where natural incubation is unlikely to be successful (e.g., because of threat of erosion, predation or collection by people), a common strategy is to move eggs to areas that can be protected until hatchlings are produced (Garcia et al. 2003). For sea turtles eggs relocated for conservation purposes, eggs should be incubated in conditions similar to those of natural nests (Mutalib and Fadzly, 2015), in order to minimize potential impacts, including those on hatchling performance (Maulany et al. 2012).

The most common nesting species of sea turtle in Pacific Guatemala is the olive ridley *(Lepidochelys olivacea)*, and egg relocation is universally used for all known sea turtle nests found on beaches along the Pacific coast. Since the early 1970s, the authorities established a conservation system using protected hatcheries, in which sea turtle eggs are reburied for incubation. Under this conservation system, egg collectors deliver 20% of the eggs from each nest and in exchange, they are allowed to sell the rest of the eggs (CONAP, 2018), either for consumption at a local marketplace or for sale to hatcheries where the excess eggs will be incubated. This study aimed to evaluate the performance of hatchling turtles, in relation to various conditions measured at the Hatchery of Multiple Uses Area of El Hawaii (AUMH), during the nesting season of 2019.

## Materials and Methods

The research was executed under Guatemalan research license number DRSO 001/2019, and data consisted in olive ridley (*L. olivacea*) hatchlings produced from eggs obtained from nests collected from the east coast of Pacific Guatemala, specifically in the area surrounding AUMH. More than 60 nighttime patrols were conducted along the beaches from Monterrico to El Hawaii, during July and August of 2019, to encounter nesting female olive ridley turtles. Once a nesting female was found, we took photos of the turtle for later assessment of asymmetry (see below), and after oviposition, we carefully removed the eggs from the nest cavity, and noted depth and width in order to replicate those conditions in the hatchery. Eggs from each nest were relocated in the hatchery, and placed in a constructed nest cavity with similar dimensions as the original nest. We also inserted a HOBO®Pendant datalogger in the middle of the egg clutch, to record temperatures every hour during the entire incubation period. After 40 days of incubation, a metal mesh cylinder was placed over the nest to retain emergent hatchlings so they could be linked to a specific nest.

Five nests were intended initially, nevertheless, data of the hatchlings (DIx and morphometric data) was able to measure at the time of their emergence, and so performance information of these hatchlings was obtained, this nest was a regular nest from the hatchery and therefore no natural nest information and temperatures were recoded. From the six nests, we assessed 210 hatchlings. The amount of hatchlings measured from each nest corresponded to all the hatchlings that emerged at the same time, thus, the amount measured hatchlings varies from 10 to 55. They were put aside and started measuring one by one. When the remaining hatchlings of each nest emerged (from 12 to 24 hours after the first group emerged), they were released as part of the usual management process, by the park rangers of volunteers.

For each hatchling assessed, we collected morphometric data including mass, with a digital scale, curved carapace length (CCL) and curved carapace width (CCW) with a flexible measuring tape, and straight carapace length (SCL) with calipers. The lengths of both front flippers were measured with a measuring tape. We also took an overhead photo of the hatchling, to calculate asymmetry based on the developmental instability index –DIx- (see below).

For performance two parameters were used: time of self righting and time of crawling from one point to another, based on the methodologies proposed by Sim et al. (2014b). Each hatchling was measured first for the self righting test, in which the hatchling was placed in a recumbent position on a surface of smoothed sand, and we recorded the time needed for the turtle to return to an upright position. For the cases when the hatchling did not successfully right itself within 10 seconds, the hatchling was manually placed in its normal position until it became active again (generally within one or 12 seconds). Subsequently, it was tested again for self righting. If a turtle failed to self-right during three trials, it was not tested again.

After the self righting test, crawling performance was measured using a “race track,” which was a 50 cm long plastic gutter filled with moist, flattened, and compact beach sand, and placed at a flat angle. Each hatchling was placed at one end and we recorded the time taken for the turtle reach the other end of the gutter, all hatchlings were measured one time in which the end was reached. Both performance measurements were made as soon as the hatchlings started to emerge from the nests (whatever hour was that happened, mainly after midnight), with data collected from one hatchlings at a time, while the other turtles were kept in an open bin apart from the measuring area. Hatchlings were retained for performance measurements no more than four hours after emergence from the nest. After the performance tests were completed, pictures were taken and morphometric measurements were collected, and subsequently all hatchlings were released into the sea. All tests were conducted in the presence of red light, to allow data collection while avoiding impacts to turtle behavior. Unfortunately, the first group of hatchlings that emerged from the fifth nest were not observed on time, and could not be used for the study, although, ten hatchlings form another emerging group of this nest were measured as soon as emerged, (the next day after the first group emerged). This lack of hatchlings was compensated with the measurement of 43 hatchling of a sixth nest.

After 4 to 6 days of the first emergence, the hatching success of each nest was based on inventory of the nest six days after first hatchling emergence, when the contents of the nest were characterized (Mutalib and Fadzly, 2015). The remains of each nest were classified as number of shells, number of hatchlings found alive within the nest, number of dead hatchlings in the nest, number of unhatched eggs, including those without apparent development and those with unhatched term embryos. Calculation of hatching success followed Miller (2000).

### Data analysis

To estimate the asymmetry of the carapace of both hatchling and mother, as part of the developmental instability, the DIx was used. This index, rather than size, contemplates relative proportions, which it achieves by integrating the analysis of the diversity of scutes by means of an averaged geometric analysis of Shannon H entropy and the difference in the costal scutes of both sides obtained through Edward’s angular distance analysis (Cortés-Gómes et al., 2018). To do this, using photographs, the costal scutes of the left and the right side were counted and measured. In order to obtain the inter-side differences, the shields of both sides were counted, while to obtain the intra-side differences. Inkscape Software 0.92.4 was used to measure each scale with a ruler. The measurements were tabulated in the program Microsoft Excel® and analyzed with the HelpersMG package (≥ 1.7) and DIx, in the Software statistical package R 4.0.2.

Using the Software statistical package R 4.0.2, we used General Linear Models (GLM) to analyze the interactions among incubation conditions, hatchling and maternal development instability, and hatchling sizes (IDx, IDxMother, minT, maxT, MeanT, MeanF, LF, RF, Eggs, HatchSuccess, NDepth, Mass, SCL, CCL, and CCW, where T = temperature, F=flipper length), and the performance of the hatchlings, in terms of times of self righting and crawling.

## Results

Nesting data, success and hatchling measurements (Table 1) were obtained from relocated nests in the hatchery of the AUMH. Complete data from nest 1 to 5 is available, unlike nest 6 which lacks data for DIx of the mother (DIxMother), number of eggs (Eggs) and nest depth (NDepth), temperatures (High, Low, Mean), and hatching success of the nest). Therefore, analyses for DIxMother, NDepth, Eggs, temperatures (High, Low, Mean), and Hatching success was for nests 1-5 only (Table 2 and Table 3).

**Table 1.** Data of the hatchlings measured from nests relocated to the hatchery of the AUMH.

| Nest | Total depth of the nest | Number of eggs | Hatching success of the nest | Number of turtles assessed | Mean crawling time | Mean self righting time | Mean Mass | Mean CCL | Mean CCW | Mean SCL |
| --- | --- | --- | --- | --- | --- | --- | --- | --- | --- | --- |
| 1 | 26 cm | 83 | 89.2 % | 34 | 103.8 s | 6.82 s | 0.019 | 4.42 | 4.45 | 4.19 |
| 2 | 29 cm | 78 | 93.6 % | 20 | 96 s | 3.12 s | 0.017 | 4.09 | 4.26 | 3.99 |
| 3 | 35 cm | 112 | 64.3 % | 55 | 122.4 s | 4.64 s | 0.017 | 4.37 | 4.31 | 4.15 |
| 4 | 34 cm | 95 | 66.7 % | 48 | 132.6 s | 6.85 s | 0.016 | 4.25 | 4.16 | 4.1 |
| 5 | 33 cm | 112 | 59.8 % | 10 | 146.4 s | 9.76 s | 0.015 | 4.46 | 4.28 | 4.18 |
| 6 | NA | NA | NA | 43 | 109.8 s | 3.53 s | 0.017 | 4.34 | 4.40 | 4.32 |

**Table 2.**
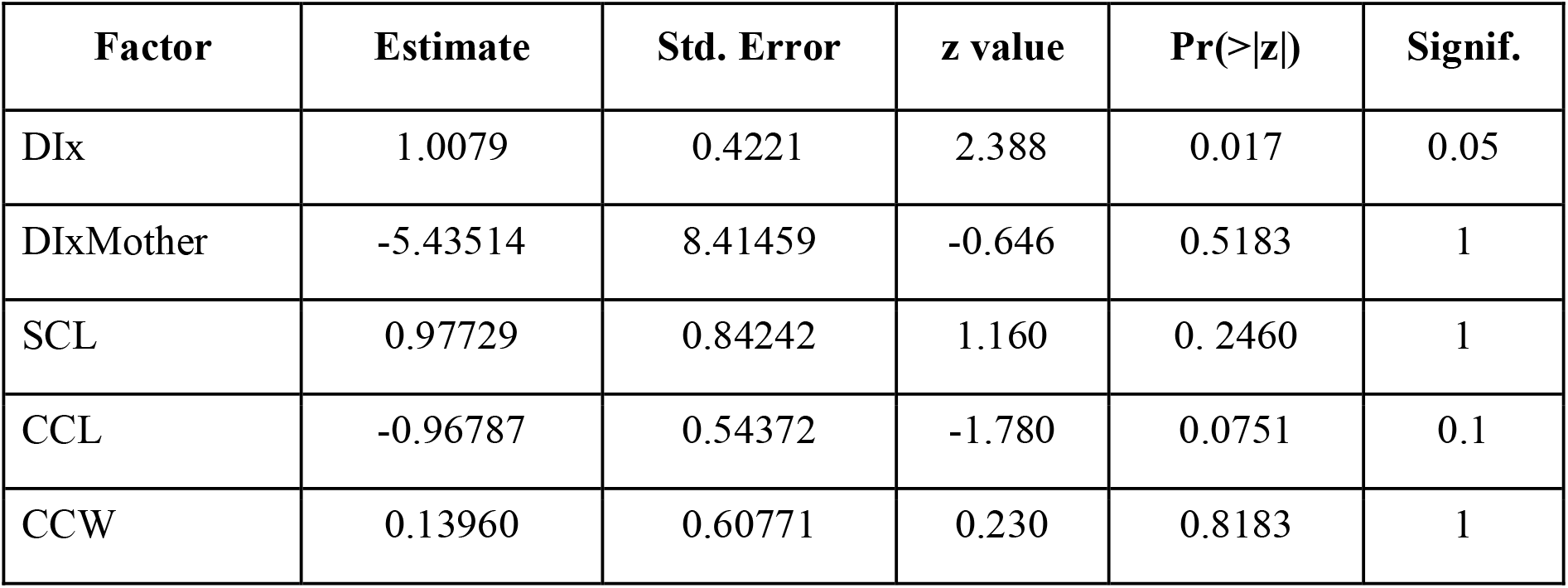
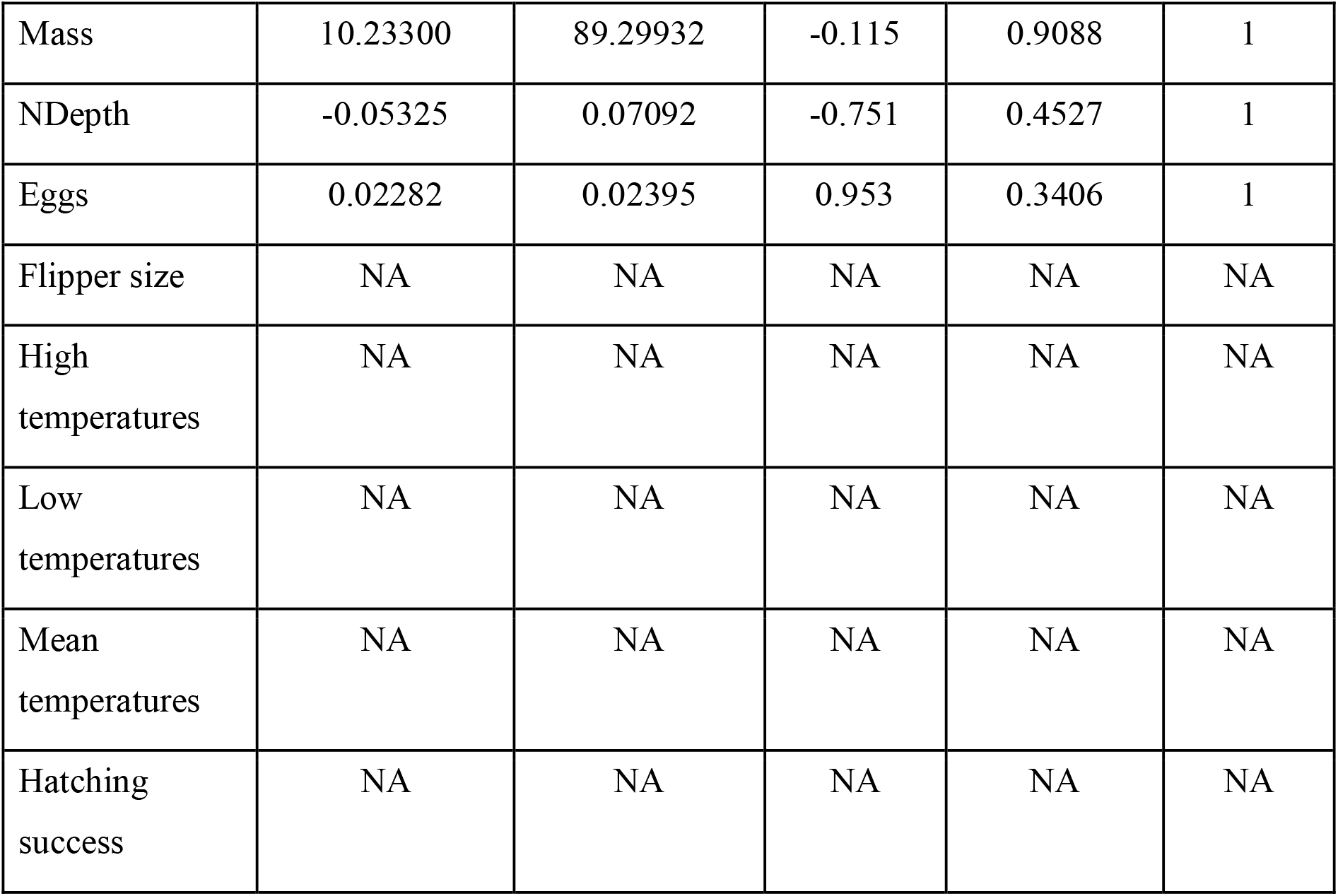
GLM results of the crawling performance and intrinsic and extrinsic factors.

| Factor | Estimate | Std. Error | z value | Pr(>\|z\|) | Signif. |
| --- | --- | --- | --- | --- | --- |
| DIx | 1.0079 | 0.4221 | 2.388 | 0.017 | 0.05 |
| DIxMother | -5.43514 | 8.41459 | -0.646 | 0.5183 | 1 |
| SCL | 0.97729 | 0.84242 | 1.160 | 0.2460 | 1 |
| CCL | -0.96787 | 0.54372 | -1.780 | 0.0751 | 0.1 |
| CCW | 0.13960 | 0.60771 | 0.230 | 0.8183 | 1 |
| Mass | 10.23300 | 89.29932 | -0.115 | 0.9088 | 1 |
| NDepth | -0.05325 | 0.07092 | -0.751 | 0.4527 | 1 |
| Eggs | 0.02282 | 0.02395 | 0.953 | 0.3406 | 1 |
| Flipper size | NA | NA | NA | NA | NA |
| High temperatures | NA | NA | NA | NA | NA |
| Low temperatures | NA | NA | NA | NA | NA |
| Mean temperatures | NA | NA | NA | NA | NA |
| Hatching success | NA | NA | NA | NA | NA |

**Table 3.**
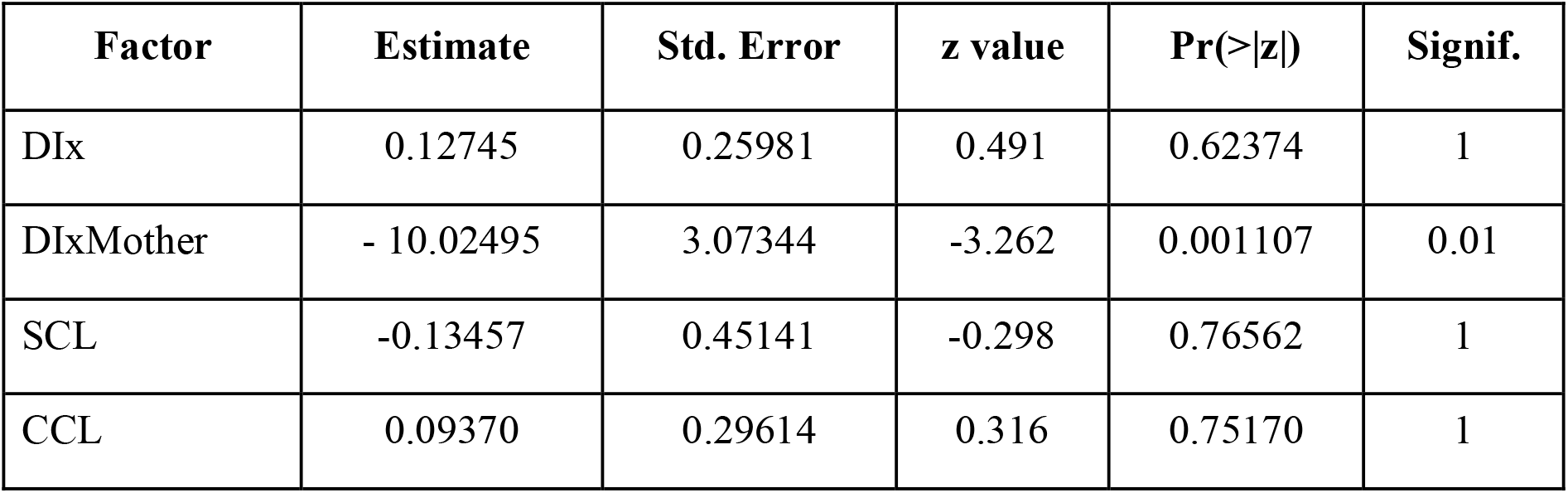
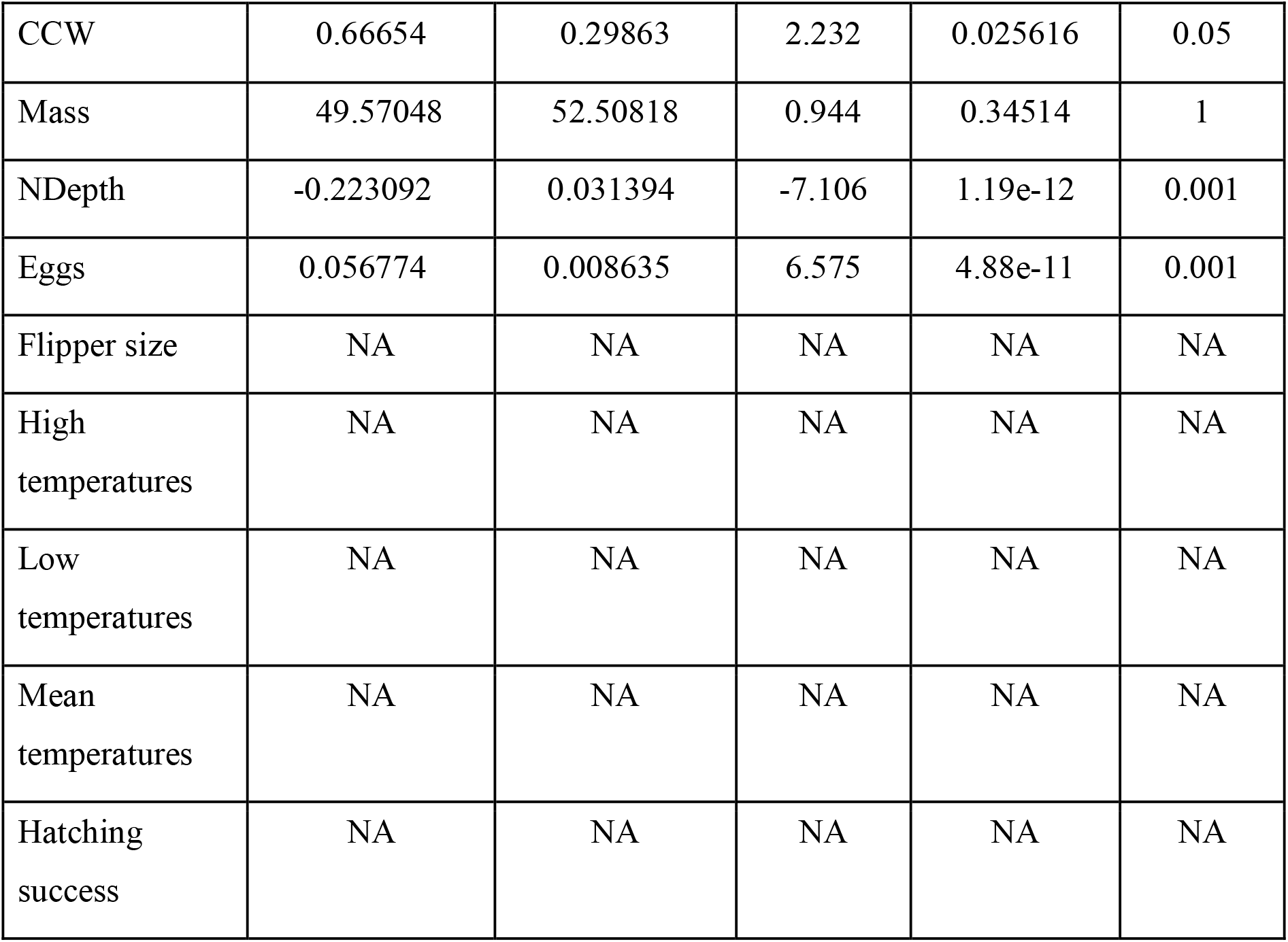
GLM results of the self righting performance and intrinsic and extrinsic factors.

| **Factor** | **Estimate** | **Std. Error** | **z value** | **Pr(>\|z\|)** | **Signif.** |
| --- | --- | --- | --- | --- | --- |
| DIx | 0.12745 | 0.25981 | 0.491 | 0.62374 | 1 |
| DIxMother | - 10.02495 | 3.07344 | -3.262 | 0.001107 | 0.01 |
| SCL | -0.13457 | 0.45141 | -0.298 | 0.76562 | 1 |
| CCL | 0.09370 | 0.29614 | 0.316 | 0.75170 | 1 |
| CCW | 0.66654 | 0.29863 | 2.232 | 0.025616 | 0.05 |
| Mass | 49.57048 | 52.50818 | 0.944 | 0.34514 | 1 |
| NDepth | -0.223092 | 0.031394 | -7.106 | 1.19e-12 | 0.001 |
| Eggs | 0.056774 | 0.008635 | 6.575 | 4.88e-11 | 0.001 |
| Flipper size | NA | NA | NA | NA | NA |
| High temperatures | NA | NA | NA | NA | NA |
| Low temperatures | NA | NA | NA | NA | NA |
| Mean temperatures | NA | NA | NA | NA | NA |
| Hatching success | NA | NA | NA | NA | NA |

We collected performance measurements on 210 hatchlings overall. Intrinsic and extrinsic factors were not found strongly significant to be related to the crawling performance, as shown in Table 2, where only the hatchlings DIx has a 0.05 significance.

Of all intrinsic and extrinsic factors, only the DIx of the hatchlings had a relationship with the crawling performance (*p*=0.05), so the focus of the results and discussion is on the self righting performance data. As shown in Table 3, the nest depth and the amount of eggs in the nest are significantly related to the self righting performance. Specifically, there is an inverse relationship between the nest depth and the self righting time, whereas the number of eggs is positively related to the amount of time it took each hatchling to flip.

Significance was also found in the mother’s DIx (DIxMother), and in the size of the hatchling, in terms of the CCW, and the self righting time.

Figure 2 shows the thermal progression of nests one to five, every hour during the whole incubation period. There was no datalogger on nest six, because is a nest for which hatchlings’ morphometric and DIx data was obtained later one, therefore, no temperature information was collected.

## Discussion

The hatchlings phenotype can be a reflection of the genetic components or maternal origins (intrinsic factors), but also can be a reflection of the incubation conditions or nest effects (extrinsic factors) (Booth et al., 2013). There are different intrinsic and extrinsic factors or conditions that can affect the performance of a hatchling, as we have shown. The intrinsic factors that are commonly studied are the characteristics directly related to the biology and physiology of the hatchlings, including maternal and genetic effects. In this study, we used the DIx metric of hatchlings and mothers, hatchlings size and mass, and flipper lengths. In addition, we considered extrinsic factors, including nest depth, clutch size, nest temperature, and hatching success (Table 1).

For marine turtles, hatchling performance measures collected soon after emergence from the nest have been used as indices of fitness (Booth et al., 2013, Fisher et al., 2014, Read et al., 2012). The performance is thought to correlate with survival because it influences the length of time a hatchling will spend on the beach, exposed to land-based predators such as ghost crabs, and some nocturnal mammals and birds (Booth et al., 2013, Ischer et al., 2009, Janzen et al., 2007, Paitz et al., 2010, Pankaew and Milton, 2017, Read et al., 2012). Freedberg et al. (2004), found that conditions experienced during development can affect the self righting response of older juveniles, indicating that the environment that the embryos experience during the incubation period can have a long term effect on its phenotype.

In this study, we found that only the DIx of hatchlings (Table 2) was significantly related to the crawling speed, despite the findings of other studies. For instance, it has been shown that hatchlings from nests with a three-day-maximum temperature below 34°C could have a faster crawling, than hatchlings with nests with three-day-maximum above 34°C (Maulany et al., 2012). Ischer et al. (2009), also found that hatchlings from cooler nests were faster crawlers. In addition, Sim et al. (2015) found that that larger hatchlings tend to be produced in cooler nests and as a result are faster crawlers, which would be expected because of their longer limbs and consequent greater stride length (Ischer et al., 2009). Interestingly, Le Gouvello et al. (2020), found no relationship between crawling speed and any of the hatchling’s attributes (different body size measures), similar to the our study, although the relationship we found was not strong.

We also looked at self righting as an indicator of performance of the hatchlings (Table 3). Contrary to what we found, previous reports have found that hatchlings from high incubation temperature nests took more time to self-right than hatchlings from cooler nests (Ashmore and Janzen, 2003, Fleming, 2019, Ischer et al., 2009, Maulany et al., 2012, Read et al., 2012, Wood et al., 2014). We found no effect of incubation temperature on self righting performance of hatchlings (Table 3).

We found that factors such as the nest depth and the amount of eggs in the nest have a significant influence over the self righting performance, where a higher number of eggs increases the time of self righting, and deeper nests produce hatchlings with faster flipping response (Figure 1). This suggests that the management decisions at the hatchery can affect hatchling performance, and thus fitness, which should be explored with more data in the future. For example, greater investigation of performance measurements can be explored, including possibly increasing the distance for crawl measurements, investigating swimming, and even reversing the order types of tests administered to hatchlings. Other studies have found differences in performance data measured in the same groups of turtles. For example, Sim et al. (2015), looking at *C. caretta*, found that the self righting performance was not informative while crawling performance was. Additionally, Gatto and Reina (2020b), using *L. olivacea*, found a positive relationship between the crawling performance and the self righting performance, although they also reported species-specific differences in performance measurements.

**Figure 1.**
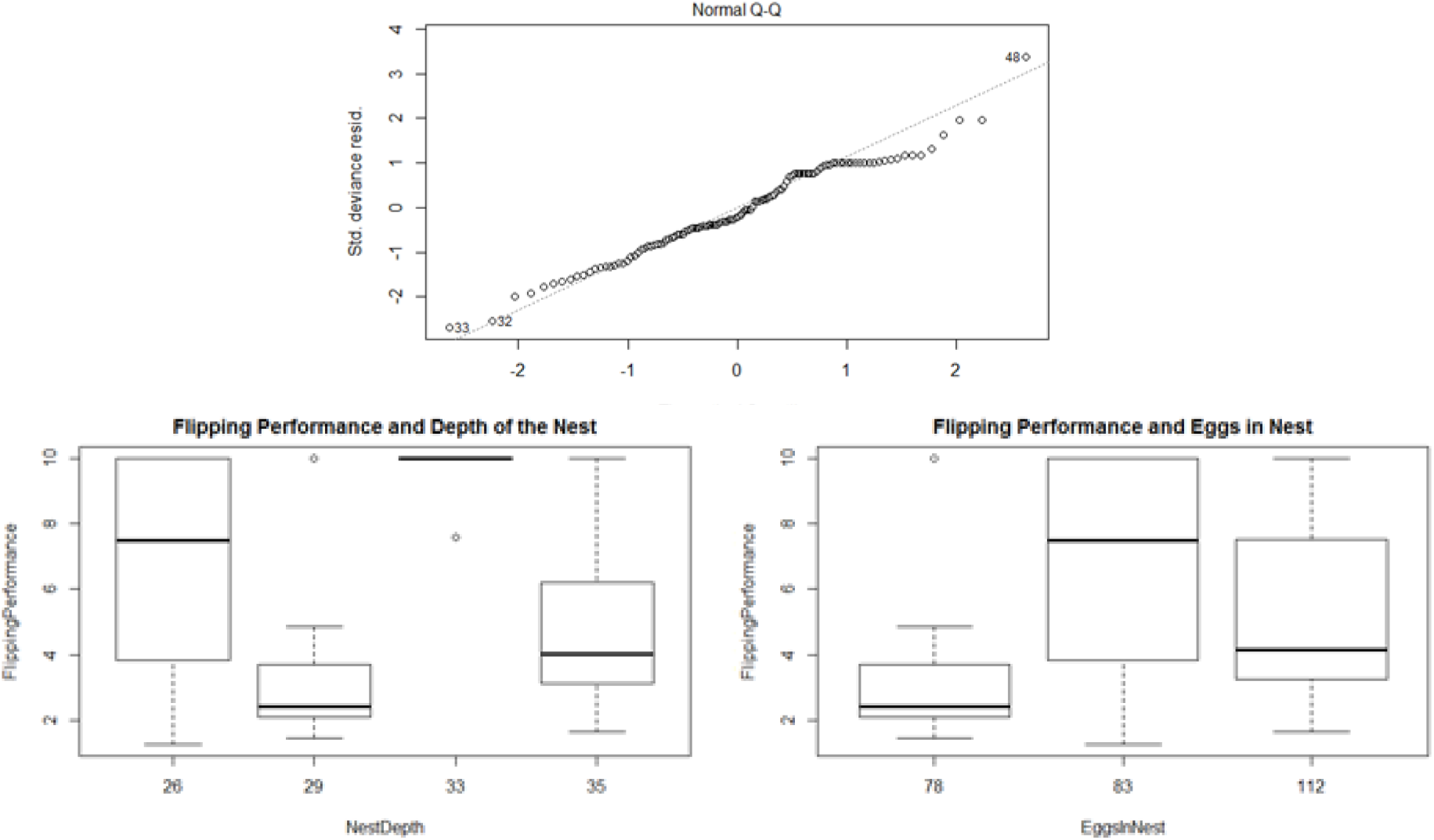
Influence of the nest depth and the number of eggs in the nest had a significant influence over the self righting performance of the 210 hatchlings measured.

There have been studies that reported how hatchlings that had incubation temperatures above 30°C needed longer time to self righting than those incubated at lower temperatures (Fisher et al., 2014, Fleming, 2019, Maulany et al., 2012). We had no hatchlings produced at incubation temperatures below 30°C, which may have obscured a significant relationship between incubation temperature and performance in this sea turtle population. Moreover, in our study, all nest temperatures at some point surpassed the upper limit of embryo tolerance (34°C), with values of 35.8-36.4°C towards the end of the incubation period (Figure 2). During the development of the embryos, when almost fully developed, high temperatures can cause uncoordinated movement in emergent hatchlings (Sim et al., 2015). Future studies should include hatchlings produced at a wider range of incubation temperatures, to better understand the relationship between incubation conditions and hatchling performance.

**Figure 2.**
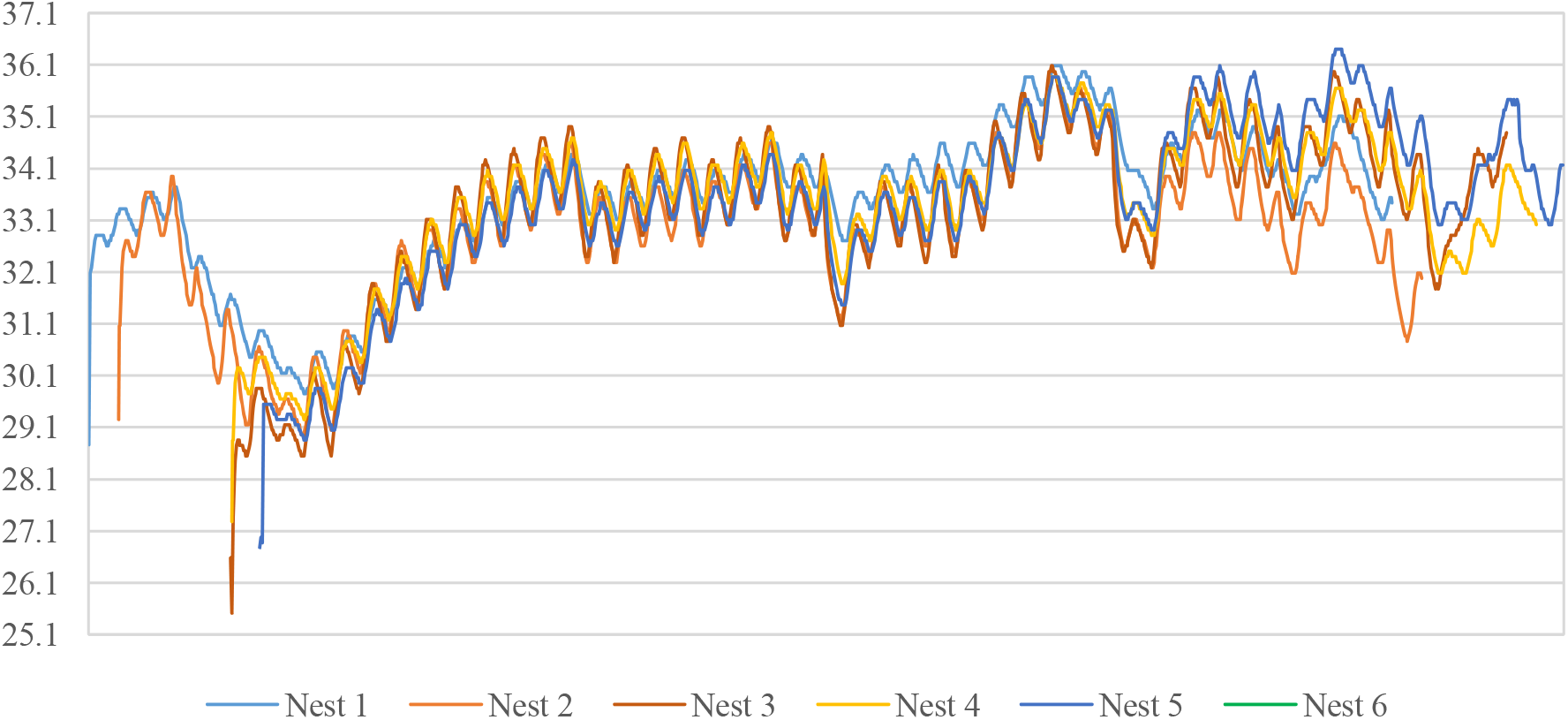
Clutch temperature, recorded every hour, in each nest of the 6 nests monitored. Starting at the nest burial until the first hatchling emerged. Note initial cold-pulse on first day when eggs were relocated to the hatchery.

Studies propose that incubation temperatures induce differences in muscle fiber composition and number within limb muscles which in turn affects locomotion performance (Ischer et al., 2009). Hatchlings that are upside down on the beach and cannot successfully self right, are more likely to die from predation, dehydration, or desiccation (Sim, 2014). Although not measured in this study, it has been reported that hatchlings use their head and neck to lift the carapace off the sand to flip-over (Read et al., 2012, Sim, 2014), which would be an interesting to study in order to understand the self righting time in hatchlings. Marine turtles have two stable balance points: the carapace and the plastron (Domokos and Várkonyi, 2008). This creates a high energy barrier between the two stable equilibria, which has to be overcome by primary biomechanical forces resulting from vertical pushing with the head against the substrate with the hyperextension of the neck (Domokos and Várkonyi, 2008). This can make neck length an important trait to consider in self righting studies (Read et al., 2012, Sim, 2014).

Other studies have reported a negative correlation between carapace size and hatchling righting time (Booth et al., 2013), with larger hatchlings reported to have a greater chance of survival because they express greater locomotor ability and ability to escape from gap-limited predators (Booth et al., 2013, Janzen et al., 2000). However, warmer incubation temperatures produce hatchlings that are smaller in size, due to a shortened period of incubation in which they have less time to convert yolk into tissue before hatching from the egg (Stewart et al., 2019). Based on this, it is possible that there is a relationship between the amount of yolk metabolized during a short incubation period and the self righting time of the hatchling.

We our data on performance of hatchlings as related to a variety of intrinsic and extrinsic incubation variables contribute to a growing number of studies on this subject (Booth 2018). We found that there is a wide range of approaches and measures used in various studies, which makes comparisons more challenging. We recommend that future studies consider following standardized approaches, including protocols for testing, analyzing and presenting data. This would greatly help facilitate comparisons across studies and improve our understanding of the impacts of incubation conditions on sea turtle fitness.

The initial objective of this study was to analyze the performance of the hatchlings in relation to the different environmental (extrinsic) and biological (intrinsic) factors experienced by developing olive ridley embryos in the hatchery environment that has been used for sea turtle egg incubation for over 50 years. However, it is likely that the hatchery conditions were not variable enough to illuminate all the potential relationships among the measures we collected. Nevertheless, based on the data collected, it appears that nest depth and number of eggs in each nest have some influence on hatchling performance, and deserve more study, in the context of informing management decisions that may improve hatchling fitness. We also suggest that some olive ridley nests are allowed to remain in place for natural incubation, in order to facilitate comparisons to hatchery-based incubation.

## Conclusions

We provide the results of the first study to assess the impacts of incubation condition on hatchling performance for olive ridley sea turtles in Guatemala. We found that self righting provided a more meaningful measure of variable behavior in response to different incubation variables than crawling speed. However, we also found there was limited variation in some incubation variables, such as temperature, due to the homogeneous environment of the hatcheries. Our results suggest that nest depth and amount of eggs in each nest can have an effect on self righting of hatchlings, with greater number of eggs result in faster flipping, while deeper nests produce slower rates of flipping by hatchlings. We recommend more studies be done on crawling performance, perhaps with a wider variety of incubation conditions and possibly longer runways, and we recognize that standardized protocols for hatchling fitness studies would facilitate better comparisons of datasets across studies.

## Competing interests

The authors received no funding for this work, as well as declare that there are no competing interests.

## Acknowledgements

We thank the Multiple Uses Area of Hawaii and ARCAS for supporting the realization of this research, as well as the volunteers that help with the data gathering. We especially would like to thank Alex García, Ricardo Gill, José Jorge Ubico, Salomé Hernández, Marisette Quiñónez, Doña Mayra, and Allan Marroquín for all their valuable help on site. Finally, we would also like to acknowledge that this research was executed under the permit no. DRSO 001-2019, granted by the Guatemalan National Council of Protected Areas (CONAP).

## Notes

### Competing Interest Statement

The authors have declared no competing interest.

